# Interval Constraint Satisfaction and Optimization for Biological Homeostasis and Multistationarity

**DOI:** 10.1101/2020.05.14.095315

**Authors:** Aurélien Desoeuvres, Gilles Trombettoni, Ovidiu Radulescu

## Abstract

*Homeostasis* occurs in a biological or chemical system when some output variable remains approximately constant as one or several input parameters change over some intervals. We propose in this paper a new computational method based on interval techniques to find species in biochemical systems that verify homeostasis. A somehow dual and equally important property is *multistationarity*, which means that the system has multiple steady states and possible outputs, at constant parameters. We also propose an interval method for testing multistationarity. We have tested homeostasis, absolute concentration robustness and multistationarity on a large collection of biochemical models from the Biomodels and DOCSS databases. The codes used in this paper are publicly available at: https://github.com/Glawal/IbexHomeo.

## 1 Introduction

The 19^th^ century French physiologist Claude Bernard, introduced the concept of homeostasis that plays a crucial role in understanding the functioning of living organisms. As he put it, homeostasis, defined as constancy, despite external changes, of the “milieu intérieur” that contains organs, tissues and cells, is a prerequisite of life. A simple example of homeostasis is the constancy of body temperature: our body temperature is maintained in a narrow range around 37°C despite large variation of the environment temperature. Another example is the concentration of many biochemical species (cell processes drivers and regulators such as glucose, ATP, calcium, potassium, cell surface receptors, transcription factors, etc.) whose steady state values are kept constant by tight control. Rather generally, homeostasis refers to constancy of the output w.r.t. variation of parameters or inputs [Golubitsky and Stewart, 2017]. Several other concepts such as robustness, resilience or viability are closely related to homeostasis and sometimes used with overlapping meaning. Robustness refers to the lack of sensitivity of temporal and static properties of systems w.r.t. parameters and/or initial conditions variation, thus encompassing homeostasis [Gorban and Radulescu, 2007; Rizk et al., 2009; Barr et al., 2019]. Resilience or viability has a more global, dynamical significance, meaning the capacity of systems to recover from perturbations via transient states that stay within bounds [Aubin, 2009].

In [Shinar and Feinberg, 2010], a special type of homeostasis is studied, named absolute concentration robustness (ACR), consisting in invariance of the steady state w.r.t. changes of initial conditions. For chemical reaction network (CRN) models, they proposed a sufficient graph-theoretical criterion for ACR. Replacing invariance by infinitesimal sensitivity, [Golubitsky and Stewart, 2017] presents a way to detect homeostasis in parametric systems of ordinary differential equations (ODEs). Their approach, based on the singularity theory, was applied to gene circuit models [Antoneli et al., 2018]. Our work builds on this reasoning, using a slightly different definition of homeostasis. Instead of looking at the infinitesimal variation, we choose to look at intervals. The steady states of ODE models are computed as solutions of algebraic equations. We are interested in finding intervals containing all the steady state values of model variables. If the variables values are contained inside sufficiently narrow intervals, those variables are stated homeostatic. Sensitivity-like calculations of the input-output relationship compute derivatives of the output w.r.t. the input and boil down to linearizations. In contrast, our method guarantees intervals containing the output w.r.t the initial system. This is crucial in applications, whenever parameters change on wide ranges and models are strongly nonlinear. The interval approach has thus larger applicability.

Our approach also provides a novel method for testing the multistationarity of biochemical models, occurring when one or several variables can have several values at the steady state. In this case, we are interested in the number and the actual space position of all the steady state solutions. Multistationarity is an important problem in mathematical biology and considerable effort has been devoted to its study, with a variety of methods: numerical, such as homotopy continuation [Sommese and Wampler, 2005] or symbolic, such as real triangularization and cylindrical algebraic decomposition [Bradford et al., 2020]. However, as discussed in [Bradford et al., 2020], numerical errors in homotopy based methods may lead to failure in the identification of the correct number of steady states, whereas symbolic methods have a double exponential complexity in the number of variables and parameters. As solving a system of algebraic equations is the same as finding intersections of manifolds, each manifold corresponding to an equation, our problem is equivalent to solving a system of constraints.

In this paper we use the interval constraint programming and optimization solvers provided by the Ibex (Interval Based EXplorer) tool, for testing homeostasis and multistationarity. Although interval solvers are combinatorial in the worst-case, polynomial-time acceleration algorithms embedded in these solvers generally make them tractable for small or medium-sized systems. Interval constraint satisfaction methods offer an interesting compromise between good precision and low complexity calculations. Interval constraint programming is an important field in computer science and its interaction with biology has cross-fertilization potential. Although interval methods have already been used in systems biology for coping with parametric models uncertainty [Tucker et al., 2007; Markov, 2010], to the best of our knowledge, this is their first application to homeostasis and multistationarity. In an algorithmic point of view, this paper reports the first (portfolio) distributed variant of the `IbexOpt` Branch and Bound optimizer, where several variants of the solver are run on different threads and exchange information. Our approach has been tested on two bio-chemical databases, and this benchmarking represents the first systematic study of homeostasis, in particular ACR, on realistic biochemical network models.

Together, our tools can be used to address numerous problems in fundamental biology and in medicine, whenever the stability and the controlability of biochemical variables are concerned. In fundamental biology, both homeostasis and multistationarity are key concepts for understanding cell decision making in development and adaptation. In personalized medicine, our tests combined with quantitative models could be used not only for a better understanding of the loss of homeostasis, for instance in aging and degenerative disease, but also for diagnosis and for predicting the effect of therapy on the re-establishment of a normal functioning.

## 2 Settings and definitions

Our definition of homeostasis is quite general and can be applied to any system of ODEs. In the applications discussed in this paper, the ODEs systems are induced by chemical reactions whose rates are given either by mass action (as in Feinberg/Shinar CRNs) or by more general kinetic laws. We consider thus a set of species variables *x*_1_, …, *x_n_*, representing species concentrations, a set of parameters *p*_1_, …, *p_r_*, representing kinetic constants, and a set of differential equations :

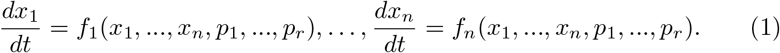

We are interested in systems that have steady states, i.e. such that the system

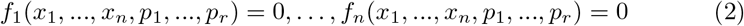

admits real solutions for fixed parameters *p*_1_, …, *p_r_*. For biochemical models, *x*_1_, …, *x_n_* represent concentrations, and in this case we constrain our study to real positive solutions.

Generally, it is possible to have one or several steady states, or no steady state at all. The number of steady states can change at bifurcations. For practically all biochemical models, the functions *f*_1_, …, *f_n_* are rational, and at fixed parameters (2) defines an algebraic variety. The local dimension of this variety is given by the rank defect of the Jacobian matrix ***J***, of elements 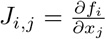, 1≤ *i*, *j* ≤ *n*.

When ***J*** has full rank, then by the implicit function theorem, the steady states are isolated points (zero dimensional variety) and all the species are locally expressible as functions of the parameters:

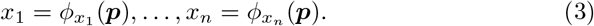

The functions *ϕ_y_* were called input-output functions in [Golubitsky and Stewart, 2017], where the input is the parameters *p_i_* and the output variable *y* is any of the variables *x_i_,* 1 ≤ *i* ≤ *n*. Also in the full rank case, a system is called *multistationary* when, for fixed parameters there are multiple solutions of (2), i.e. multiple steady states.

***J*** has not full rank in two cases. The first case is at bifurcations, when the system output changes qualitatively and there is no homeostasis. The second case is when (1) has *l ≤ n* independent first integrals, i.e. functions of *x* that are constant on any solution of the ODEs (1). In this case the Jacobian matrix has rank defect *l* everywhere and steady states form a *l*-dimensional variety. For instance, for many biochemical models, there is a full rank constant matrix *C* such that 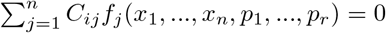, for all *x*_1_, …, *x_n_, p*_1_, …, *p_r_,* 1 ≤ *i* ≤ *l*. In this case there are *l* linear conservation laws, i.e. 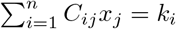, 1 ≤ *i* ≤ *l* are constant on any solution. Here *k_i_* depends only on the initial conditions, 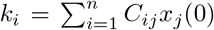. In biochemistry, linear conservation laws occur typically when certain molecules are only modified, or complexified, or translocated from one compartment to another one, but neither synthesized, nor degraded. The constant quantities *k_i_* correspond to total amounts of such molecules, in various locations, in various complexes or with various modifications.

A biological system is characterized not only by its parameters but also by the initial conditions. For instance, in cellular biology, linear conservation laws represent total amounts of proteins of a given type and of their modifications, that are constant within a cell type, but may vary from one cell type to another. Therefore we are interested in the dependence of steady states on initial conditions, represented as values of linear conservation laws. Because conservation laws can couple many species, steady states are generically very sensitive to their values. ACR represents a remarkable exception when steady states do not depend on conservation laws. In order to compute steady states at fixed initial conditions, we solve the extended system

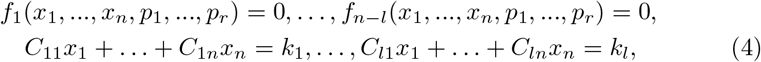

where *k_i_* are considered as extra parameters, and *f*_1_, …, *f_n_*_−*l*_ are linearly independent functions. In this case, excepting the degenerate steady states with zero concentrations discussed at the end of this section, the Jacobian of the extended system has full rank and one can define again input-output functions as unique solutions of (4). A system is *multistationary* if at fixed parameters there are multiple solutions of (4).

*Homeostasis* is defined using the input-output functions.

### Definition 21.

*We say that y is a k-homeostatic variable if in the path of steady states given by Φ_y_ we get :*

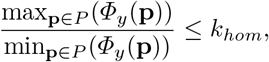

where *k_hom_* ≥ 1 is a small positive real number. We take *k_hom_* = 2 by default, but we can choose a smaller *k_hom_* if the parameter variation is small. *P* represents the space of parameters (we take a *P* compact for our examples), and **p** is a point inside *P* . So, we consider homeostasis of a variable *y* for any change of parameters in a domain *P*.

We exclude from our definition trivial solutions *x_i_* = 0 obtained when

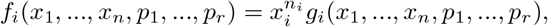

where *n_i_* are strictly positive integers, *g_i_* are smooth functions with non-zero derivatives *∂f_i_/∂x_i_* for *x_i_* = 0. These solutions persist for all values of the parameters and are thus trivially robust. In this case we replace the problem *f_i_* = 0 by the problem *g_i_* = 0 that has only non-trivial solutions *x_i_* ≠ 0.

## 3 Interval methods for nonlinear constraint solving and optimization

### 3.1 Intervals

Contrary to standard numerical analysis methods that work with single values, interval methods can manage sets of values enclosed in intervals. By these methods one can handle exhaustively the set of possible constraint systems solutions, with guarantees on the answer. Interval methods are therefore particularly useful for handling nonlinear, non-convex constraint systems.

#### Definition 31.

*An* interval 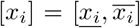 defines the set of reals *x_i_* such that 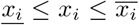. 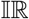 denotes the set of all intervals. A **box** [*x*] denotes a Cartesian product of intervals [*x*] = [*x*_1_] × … × [*x_n_*]. The size or width of a box [*x*] is given by *w*[*x*] ≡ max_*i*_(*w*([*x_i_*])) where 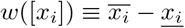.

*Interval arithmetic* [Moore, 1966] has been defined to extend to 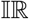 the usual mathematical operators over ℝ (such as +, ., */*, power, sqrt, exp, log, sine). For instance, the interval sum is defined by 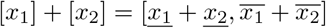. When a function *f* is a composition of elementary functions, an *extension* of *f* to intervals must be defined to ensure a conservative image computation.

#### Definition 32. (Extension of a function to 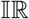)

*Consider a function f* : ℝ^*n*^ → ℝ.

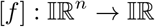 *is said to be an* **extension***of f to intervals iff:*

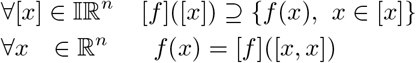

The *natural extension* of a real function *f* corresponds to the mapping of *f* to intervals using interval arithmetic. More sophisticated interval extensions have been defined, based on interval Taylor forms or exploiting function monotonicity [Jaulin et al., 2001].

### 3.2 Interval methods for constraint solving

Several interval methods have been designed to approximate *all* the real solutions of equality constraints (*h*(*x*) = 0) in a domain defined by an initial box [*x*]. These methods build a search tree that explores the search space exhaustively by subdividing [*x*]. The tree built contains a set of nodes, each of them corresponding to a sub-box of [*x*]. At each node, the Branch and Contract process achieves two main operations:

- **Bisection**: The current box is split into two sub-boxes along one variable interval.
- **Contraction**: Both sub-boxes are handled by *contraction* algorithms than can remove sub-intervals without solution at the bounds of the boxes.

At the end of this tree search, the “small” boxes of size less than a user-given precision *ϵ* contain all the solutions to the equation system. The process is combinatorial, but the contraction methods are polynomial-time acceleration algorithms that make generally the approach tractable for small or medium-sized systems. Without detailing, contraction methods are built upon interval arithmetic and can be divided into constraint programming (CP) [Van Hentenryck et al., 1997; Neveu et al., 2015; Benhamou et al., 1999] and convexification [Tawarmalani and Sahinidis, 2005; Misener and Floudas, 2014] algorithms.

## 4 Multistationarity

The constraint solving strategy roughly described above is implemented by the `IbexSolve` strategy available in the `Ibex` C++ interval library. `IbexSolve` can find all the solutions to the system (4) with fixed parameters in a straightforward way.

This method is useful for small and medium systems, and sometimes for large systems, depending on the nature of the constraints and the efficiency of the contractors. Also, it provides as output each solution box. This output is easy to read, because (4) has always a finite set of solutions.

In case of large systems, it can be easier to answer the question: do we have zero, one, or several steady states? In this case, we can use another strategy, described in the next section, where the problem is reformulated in terms of 2*n* constrained global optimization problems: for every variable *x_i_*, we call twice an optimization code that searches for the minimum and the maximum value of *x_i_* while respecting the system (4).

- If the system admits at least two distinct solutions, the criterion used in Definition 21 (using *k* close to 1) will fail for at least one species, i.e. we will find a species *x_i_* whose minimum and maximum values are not close to each other.
- If the system admits no solution, the first call to the optimizer (i.e., minimizing *x*_1_) will assert it.
- And if we have only one solution, every species will respect the criterion.

Let us give a simple example given by the model 233 in the Biomodels database. In this model we have two species *x* and *y* together with seven parameters (one for the volume of the compartment, four for kinetic rates, and two for assumed fixed species). The system of ODEs is given by:

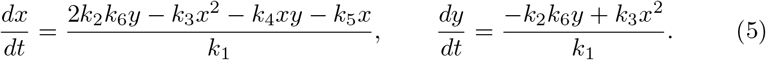

After replacing the parameters by their values, the steady state equations read:

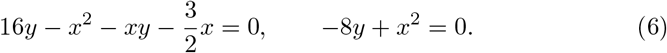

The system (6) has two non-zero solutions, given by (6,4.5) and (2,0.5). When the system (6) is tested by `IbexHomeo` (the dedicated strategy for homeostasis) on a strictly positive box (to avoid the trivial solution (0,0)), we find *x* ∈ [2, 6] and *y* ∈ [0.5, 4.5]. The homeostasy criterion fails at fixed parameters and we know that we have multistationarity.

## 5 IbexHomeo for finding homeostatic species

The new interval solver dedicated to homeostasis proposed in this paper resorts to several calls to optimization processes. Let us first recall the principles behind interval Branch and Bound codes for constrained optimization.

### 5.1 Interval Branch and Bound methods for constrained global optimization

Constrained global optimization consists in finding a vector in the domain that satisfies the constraints while minimizing an *objective function*.

#### Definition 51.

*Let x* = (*x*_1_, …, *x_n_*) *varying in a box* [*x*]*, and functions f* : ℝ*^n^* → ℝ*, g* : ℝ*^n^* → ℝ*^m^, h* : ℝ*^n^* → ℝ^*p*^.

*Given the system S* = (*f, g, h, x,* [*x*]), *the constrained global optimization problem consists in finding f*\*:

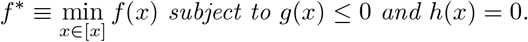

*f denotes the objective function, f*\* *being the objective function value (or best “cost”), g and h are inequality and equality constraints respectively. x is said to be feasible if it satisfies the constraints.*

Interval methods can handle constrained global optimization (minimization) problems having non-convex operators with a Branch and Bound strategy generalizing the Branch and Contract strategy described in the previous section. The Branch and Bound solver maintains two bounds *lb* and *ub* of *f*\*. The upper bound *ub* of *f*\* is the best (lowest) value of *f* (*x*) satisfying the constraints found so far, and the lower bound *lb* of *f*\* is the highest value under which it does not exist any solution (feasible point). The strategy terminates when *ub − lb* (or a relative distance) reaches a user-defined precision *ϵ_f_* . To do so, a variable *x_obj_* representing the objective function value and a constraint *x_obj_* = *f* (*x*) are first added to the system. Then a tree search is run that calls at each node a bisection procedure, a contraction procedure, but also an additional bounding procedure that aims at decreasing *ub* and increasing *lb*. Improving *lb* can be performed by the contraction procedure: it is given by the minimum value of 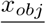 over all the nodes in the search tree. Improving the upper bound is generally achieved by local numerical methods. Like any other Branch and Bound method, improving the upper bound *ub* allows the strategy to eliminate nodes of the tree for which 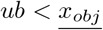.

*Remark.* Interval Branch and Bound codes can solve the optimization problem defined in Def. 51 but they sometimes require a significant CPU time because of the guarantee on the equality constraints. A way to better tackle the problem in practice is to relax equalities *h*(*x*) = 0 by pairs of inequalities −*ϵ_h_ ≤ h*(*x*) and *h*(*x*) ≤ +*ϵ_h_*, where *ϵ_h_* is a user-defined positive parameter. Therefore, in practice, interval Branch and Bound codes generally compute a feasible vector *x* satisfying the constraints *g*(*x*) ≤ 0 and −*ϵ_h_ ≤ h*(*x*) ≤ +*ϵ_h_* such that |*f*\* − *f*(*x*)| ≤ *ϵ_f_*.

The interval Branch and Bound strategy roughly described above is implemented by the `IbexOpt` strategy available in the `Ibex` C++ interval library [Chabert, 2020]. `IbexOpt` is described in more details in [Trombettoni et al., 2011; Neveu et al., 2016].

### 5.2 A dedicated solver for Homeostasis based on IbexOpt

Remember that we consider a variable *x_i_* to be homeostatic if it verifies Definition 21. For identifying homeostasis, we consider the system (4) in which the parameters *p_i_* and *k_i_* can vary.

#### Bi-optimization for a given species *x*_i_

Since we want to compute the minimum and the maximum value of *x_i_* = *Φ_x_i* (**p**), the homeostasis detection amounts to two optimization problems, one minimizing the simple objective function *x_i_*, and one maximizing *x_i_*, i.e. minimizing −*x_i_*. The two values returned are finally compared to decide the *x_i_* homeostasis. It is useful to consider that minimizing and maximizing *x_i_* are somehow symmetric, allowing the strategy to transmit bounds of *x_i_* from one optimization process to the dual one. These bounds can also be compared during optimization to stop both optimizations if they give enough information about homeostasis. Indeed, an optimizer minimizing *x_i_* computes [*ln, un*] ∋ min(*x_i_*), where *ln* and *un* are *lb* and *ub* of the objective function *x_i_*. An optimizer maximizing *x_i_* computes [*ux, lx*] ∋ max(*x_i_*), where *ux* and *lx* are −*ub* and −*lb* of the objective function −*x_i_*. Without detailing, *lx/ln* is an overestimate of the “distance” between any two feasible values of *x_i_*, and a small value states that the species is homeostatic (see Def. 21). Conversely, *ux/un* is an underestimate of any two feasible values distance, and *ux/un > k* asserts that the species is not homeostatic. This TestHom decision procedure is implemented by Algorithm 1.

**Algorithm 1:**
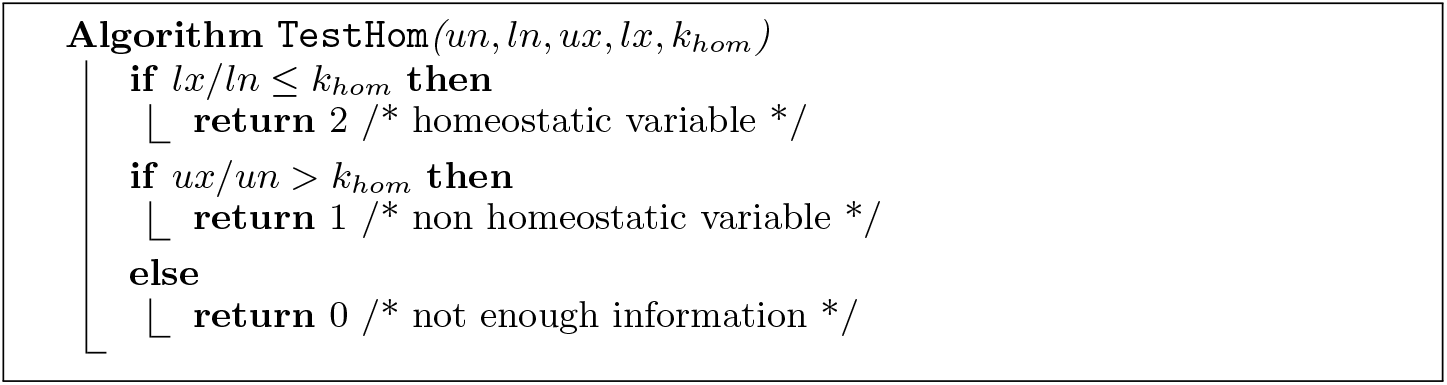
The `TestHom` decision procedure.

#### Improving upper bounding with fixed parameters

The bi-optimization described above run on the system *S* corresponding to the system (4), where the equations *f_j_*(*x, p*) = 0 are relaxed by inequalities −*ϵ_h_ ≤ f_j_*(*x, p*) ≤ +*ϵ_h_*; the parameters *p* can vary in a box [*p*] and are added to the set of processed variables. But it is important to notice that a steady state is expected for *every* parameter vector *p* ∈ [*p*] (this is not valid, for instance, in the neighborhood of a saddle-node bifurcation, which should be avoided by re-defining [*p*]). We exploit this key point by also running minimization and maximization of *x_i_* on a system *S*′, corresponding to the system *S* where the parameters have been fixed to a random value *p* ∈ [*p*], with the hope that reducing the parameter space allows a faster optimization. The computed values constitute feasible points for the initial problem (i.e., with parameters that can vary) and can fasten the bi-optimization algorithm described above. Recall indeed that finding feasible points enables to improve the upper bound *ub* of *f*\* and to remove from the search tree the nodes with a greater cost.

Overall, homeostasis detection of species *x_i_* is performed by Algorithm 2.

**Algorithm 2:**
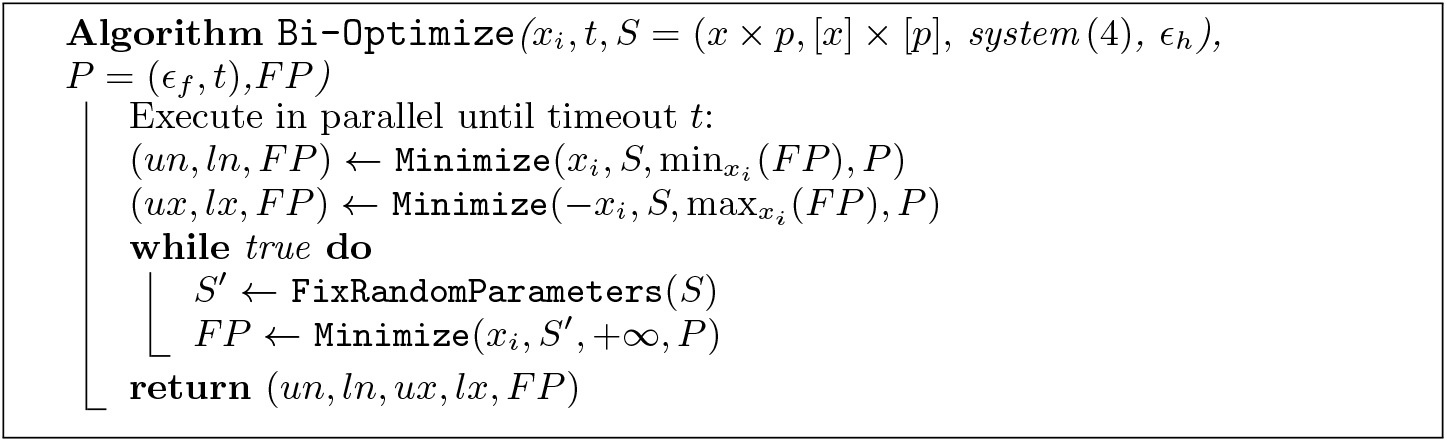
The double optimization process on a given species *x_i_*. *P* is the set of solver parameters: *ϵ_f_* is the user-defined precision on the objective function value, *t* is the timeout required.

All the optimization processes are run in parallel and exchange newly found feasible points stored in *FP* . Every call to `Minimize` on *S* can start with an initial upper bound initialized with the best feasible point found so far (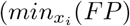 or 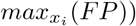).

The minimization processes on *S*′ are generally fast so that several ones can be called in a loop (with different parameters fixed to random values) until the end of the main minimization processes on *S*.

#### A portfolio strategy for the bi-optimization

It is important to understand that `IbexSolve` and `IbexOpt` are generic strategies. That is, different procedures can be selected for carrying out the choice of the next variable interval to bisect (called branching heuristic) or for selecting the next node to handle in the search tree. It is known that some heuristics in general useful can be sometimes bad for some specific problems. Therefore we propose a *portfolio* parallelization strategy where different processes (threads) run Branch and Bound algorithms using different branching heuristics (called *cutters* hereafter) or node selection heuristics (called *nodeSel*). These threads can communicate their bounds to each other, reducing the risks of an ineffective strategy. In practice, we should modify a call to `Minimize` as follows:

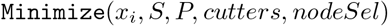

where *cutters* denotes a set of branching heuristics and *nodeSel* denotes a set of node selection heuristics. This routine calls |*cutters*| × |*nodeSel*| threads, each of them corresponding to one Branch and Bound using one branching heuristic in *cutters* and one node selection heuristic in *nodeSel*. These threads work in the same time on the same problem, but they build different search trees. Therefore one optimizer can compute an *lb* value better (greater) than the others. In this case, it sends it to the other threads.

Heuristics used to split a box are all the variants of the *smear* branching strategy described in [Trombettoni et al., 2011] and [Araya and Neveu, 2018]. Strategies used to select the next node to be handled are described in [Neveu et al., 2016]. The cutting strategy *lsmear* [Araya and Neveu, 2018] is generally more efficient than the others, and will be more often used.

#### The main IbexHomeo algorithm

Finally, because we want to determine all the homeostatic species, we run the double optimization *n* times, for every species *x_i_*, as shown in Algorithm 3. After a first call to a `FirstContraction` procedure that contracts the domain [*x*] × [*p*], `IbexHomeo` calls two successive similar loops of different performance. The first loop iterates on every species *x_i_* and calls on it the double optimization function `Bi-Optimize`. The optimization threads are all run using the *lsmear* branching heuristic and have a “short” timeout in order to not be blocked by a given species computation. If a bi-Optimization call on *x_i_* reaches the timeout *t* without enough information about homeostasis, *x_i_* is stored in *L* and the computation on subsequent species continues and can learn (and store in *FP*) new feasible points than can be exploited by other optimization processes. Indeed, the feasible region defined by *S* is the same for each optimization. Therefore the second loop is similar to the first one, but with a greater timeout and more threads in parallel running the optimization with more various branching heuristics.

To summarize, the `IbexHomeo` algorithm creates communicating threads for:

- exploiting the duality min/max of the bi-optimization related to a given species homeostasis detection,
- finding feasible points more easily,
- running a portfolio of similar Branch and Bound algorithms using different heuristics.

## 6 Experimental results

For benchmarking the multistationarity test we have used DOCSS (Database of Chemical Stability Space, http://docss.ncbs.res.in), a repository of multistationary biochemical circuits. DOCSS contains biochemical circuits with up to four species and up to five catalytic reactions. The catalytic reactions are decomposed into several mass action laws, elementary steps. In DOCSS, the models are specified as short strings of symbols coding for the catalytic reactions and as lists of numeric parameters. These specifications were first parsed to SBML files, then to systems of differential equations and conservation laws using tools developed in [Lüders et al., 2020], and transformed into an input file for our algorithms. For the benchmarking we have selected all the 210 DOCSS circuits with 3 species (denoted *a*,*b*,*c*) and 3 catalytic reactions. The mass action models have up to 6 variables (i.e., the species *a*,*b*,*c*, and several complexes resulting from the decomposition of catalytic reactions into mass action steps). The steady states of all models in DOCSS were numerically computed in [Ramakrishnan and Bhalla, 2008] using a homotopy continuation method [Sommese and Wampler, 2005]. For all the 3 × 3 models both homotopy and interval `IbexSolve` methods find 3 or 4 steady states. Although the positions of most of the solutions are almost identical using the two methods (see Figure 1), there are a few exceptions where the two solutions diverge. We have investigated each of these exceptions. The result is presented in Table 1.

**Algorithm 3:**
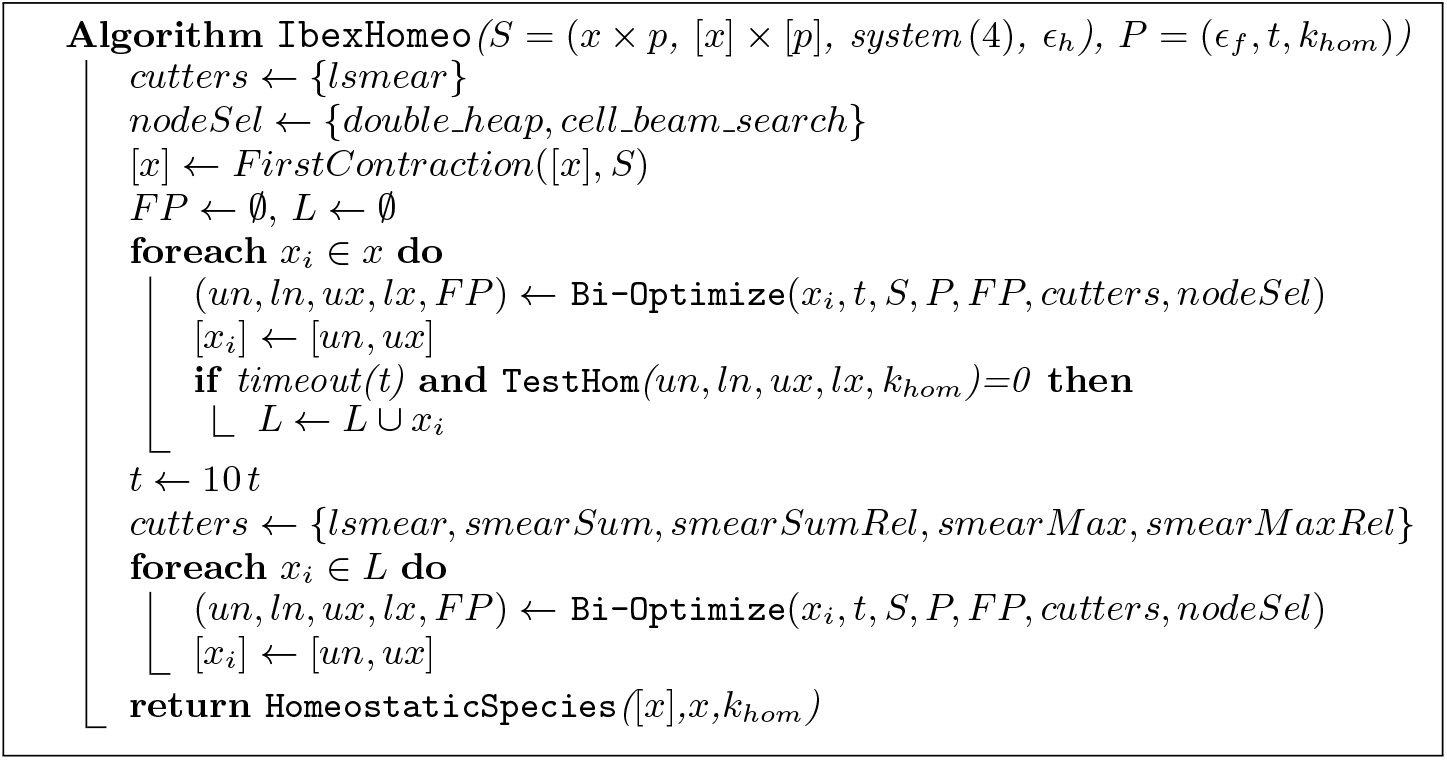
Main frame of IbexHomeo. *k_hom_* ∈ [1, 2] is defined in Def. 21. Via the procedure `HomeostaticSpecies`, the algorithm returns the set of homeostatic variables.

**Table 1.** Comparison of most divergent Ibex vs. homotopy solutions to symbolic solutions. *n* is the number of steady states. *dist* is the distance between sets of steady states solutions computed by homotopy or IbexSolve and the symbolic solutions, computed as 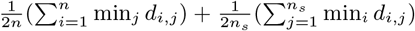, where *d_i,j_* is the Euclidean distance between the numerical solution *i* and the symbolic solution *j*, *n_s_* is either the number of solutions *n_h_* found by the homotopy method or the number *n_i_* found by IbexSolve.

| model | $n$ sym | $n_h$ homo | $n_i$ Ibex | $dist$ homo | $dist$ Ibex |
| --- | --- | --- | --- | --- | --- |
| M338-2 | 3 | 4 | 3 | 0.020087 | 2.9101e-05 |
| M464-1 | 3 | 4 | 3 | 0.012388 | 4.7049e-05 |
| M464-2 | 3 | 4 | 3 | 0.029019 | 3.5893e-05 |
| M488-1 | 3 | 4 | 3 | 0.011739 | 2.4117e-05 |
| M488-2 | 3 | 4 | 3 | 0.010935 | 2.3473e-05 |
| M488-3 | 3 | 4 | 3 | 0.017165 | 5.5298e-05 |
| M506-1 | 4 | 3 | 4 | 0.10086 | 2.6487e-05 |
| M506-2 | 4 | 3 | 4 | 0.0069613 | 4.3250e-06 |
| M506-14 | 4 | 3 | 4 | 0.0092411 | 5.0559e-05 |
| M95-1 | 4 | 3 | 4 | 0.0033133 | 1.1911e-05 |
| M95-2 | 4 | 3 | 4 | 0.00063113 | 1.1230e-05 |

**Fig. 1.**
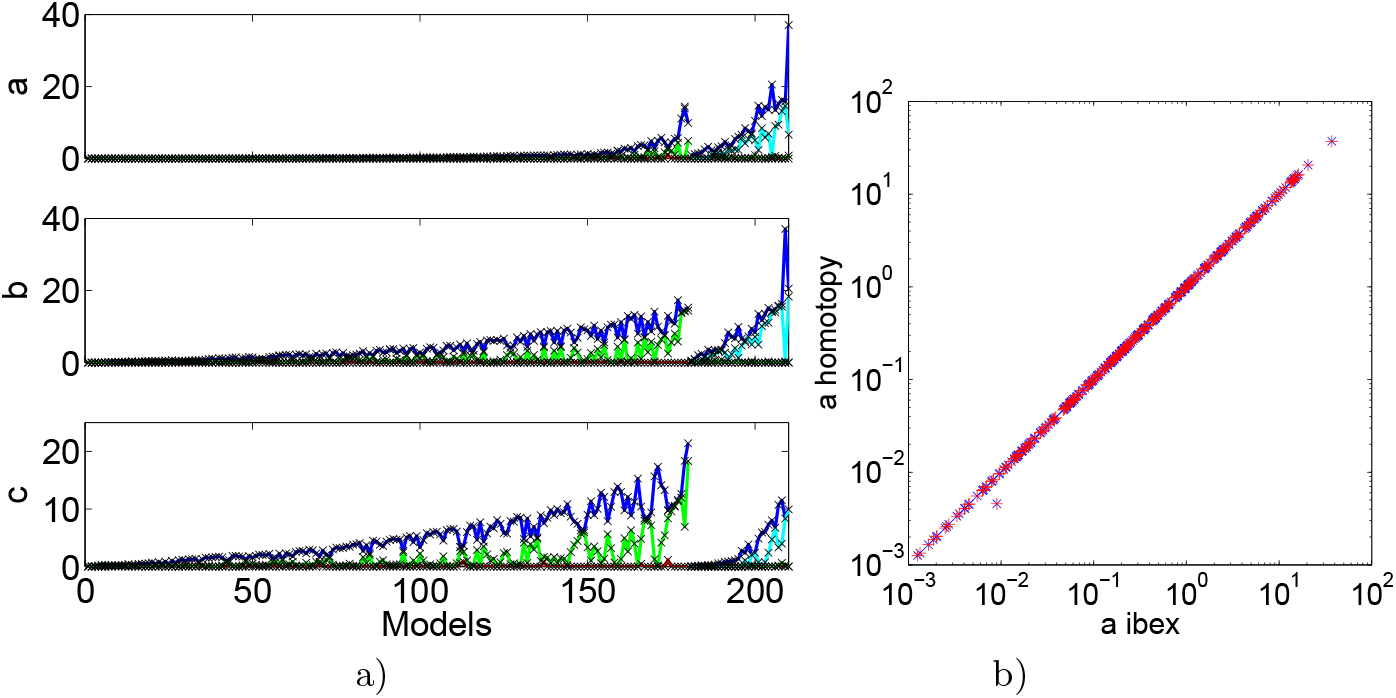
Comparison between homotopy and `IbexSolve` steady states. All the tested models are multistationary. a) Models were partitioned into two classes, with 3 (appearing first) and 4 homotopy solutions, then sorted by the average of the steady state concentrations in the homotopy solutions. Homotopy and `IbexSolve` solutions are represented as lines (red, green and blue for models with 3 steady states, cyan for the fourth) and crosses, respectively. b) Values of the steady state concentration *a* computed by homotopy and `IbexSolve`. Each `IbexSolve` steady state was related to the closest homotopy state (red +), in the Euclidean distance sense; reciprocally, each homotopy state was related to the closest `IbexSolve` state (blue crosses).

The main reason of discrepancy is a different number of solutions computed by the two methods. For the models with discrepancies we have also computed symbolic steady state solutions using the Symbolic Math Toolbox of Matlab R2013b (MathWorks, Natick, USA), though this was not possible for all the models. The comparison to `IbexSolve` and homotopy solutions shows that `IbexSolve` always finds the right number of solutions in a fraction of a second and computes their positions with better precision than the homotopy method. We conclude that discrepancies result from the failure of the homotopy method to identify the right number of solutions.

The homeostasis tests were benchmarked using the database Biomodels (https://www.ebi.ac.uk/biomodels/), a repository of mathematical models of biological and biomedical systems; parsed from SBML files to systems of differential equations and conservation laws using tools developed in [Lüders et al., 2020], and transformed into an input file for our algorithms. Of the 297 models initially considered, 72 were selected. These models have a unique steady state where every species has a non null concentration. To select them we have considered several tests described in Table 2.

**Table 2.**
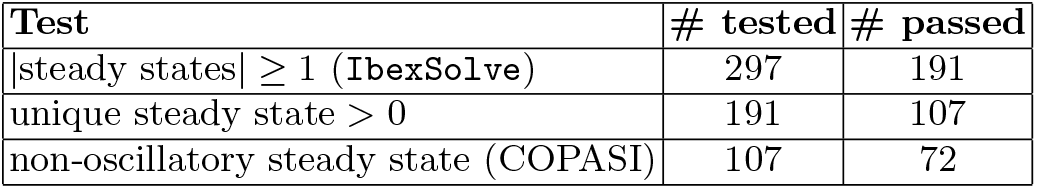
The methodology applied to select models to be tested for homeostasis from the initial set of models. A test using `IbexSolve` guarantees the existence and position of all steady states. Then, each model with a unique steady state having a non zero concentration is selected. To avoid false positive answer due to a given precision, a time course beginning from the steady state indicated by `IbexSolve` has been done using COPASI. These false positive answers may occur in a limit cycle or in a focus since we use relaxed equalities.

| Test | # tested | # passed |
| --- | --- | --- |
| \|steady states\| $\geq 1$ (**IbexSolve**) | 297 | 191 |
| unique steady state $> 0$ | 191 | 107 |
| non-oscillatory steady state (COPASI) | 107 | 72 |

The selected models have been then partitioned into three categories depending on the possible tests: kinetics rates, conservation laws, and volume compartments. We have also tested for ACR the three models described in [Shinar and Feinberg, 2010] in which the parameters have been fixed to random values. These models were previously tested for ACR by the Shinar/Feinberg topological criterion, therefore should remain so for any parameter set. As expected, the three models respect the ACR condition.

The initial boxes/domains for the conservation laws and parameters values were determined from the nominal initial conditions and parameter values found in the SBML files. The initial intervals bounds were obtained by dividing and multiplying these nominal values by a factor 10 for total amount of conservation laws and for volume compartments, and by a factor 100 for kinetics parameters, respectively. Homeostasis was tested using Definition 21. ACR was tested using Definition 21 with *k* → 1, i.e. almost zero width intervals, and where the varying input parameters are the conservation laws values.

The execution time statistics are given in Table 3.

**Table 3.** Statistics on the homeostasis test using `IbexHomeo`. Selected biomodels have been classified for three tests. All of them have been tested w.r.t. the kinetics rates, and each model presenting at least one conservation law have been tested for ACR. Moreover, models with several compartments have been tested w.r.t. their volume. As we have many timeout (computed as 360s per species, with an average dimension of 51.92 (15.48 for species, and 36.4 for parameters)), the time column consider only models that passed the test. The two last columns indicate the models with ACR in the first test, or with a 2-homeostatic species in other cases (because we get data during computation it can happen that a timeout model gives us an homeostasis).

| Test **IbexHomeo** | # models | # timeout | time (s) (min/median/max) | yes | no |
| --- | --- | --- | --- | --- | --- |
| ACR only | 33 | 17 | 0.19/4.32/10887 | 3 | 14 |
| kinetics only | 72 | 41 | 0.4/122/1923 | 4 | 29 |
| compartments only | 14 | 9 | 0.4/70/189 | 3 | 3 |

We have used Biomodels also for multistationarity tests. Among the 297 models tested for multistationarity using `IbexSolve`, 63 provide a timeout. For the solved models, 35 do not have steady-state, 153 have a unique steady state, 42 provide multistationarity, 4 have a continuum of steady states, see Appendix 8.

## 7 Discussion and conclusion

The results of the tests show that interval methods are valuable tools for studying multistationarity and homeostasis of biochemical models.

In multistationarity studies, our interval algorithms outperform homotopy continuation based numerical methods. They have found the correct number of steady states, and with a high accuracy. In terms of complexity of calculations they behave better than symbolic methods. Indeed, `IbexSolve` solve all the models in the chosen database (DOCSS). The 210 DOCSS models correspond to 13 different symbolic systems of equations (the remaining differences concern numerical parameters). The symbolic solver did not find explicit solutions for 1 of these symbolic systems, reduced 3 other models to 4^*th*^ degree equations in 3.5 to 47 s, and solved the 9 others in times from 2 to 20 s which should be compared to the fractions of a second needed for the `IbexSolve` calculations. We did not perform symbolic calculations on more complex models from Biomodels, that we expect out of reach for these methods. However, BIOMD26 was studied elsewhere [Bradford et al., 2020], and it was only with sophisticated methods that the parametric dependence of the solutions was solved symbolically for 3 parameters (with the remaining parameters being given numerical values). These results are very encouraging since multistationarity is a computationally hard problem with numerous applications to cell fate decision processes in development, cancer, tissue remodelling. As future work, we plan to test multistationarity of larger models using interval constraint programming.

In homeostasis studies, interval methods perform well for small and medium size models in the Biomodels database. However, given Claude Bernard’s statement that stability is a prerequisite of life, the result of these tests is somehow surprising. Only 18% of the tested models have some form of homeostasis. This low proportion could be explained by the possible incompleteness of the biochemical pathways models. Not only these models are not representing full cells or organisms, but they may also miss regulatory mechanisms required for homeostasis. Negative feed-back interaction is known to be the main cause of homeostasis (although feed-forward loops can also produce homeostasis) [Cooper, 2008]. As well known in machine learning, it is notoriously difficult to infer feed-back interaction. For this reason, many of the models in the Biomodels database were built with interactions that are predominantly forward and have relatively few feed-back interactions. It is therefore not a surprise that models that were on purpose reinforced in negative feed-back to convey biological homeostasis, such as BIOMD41, a model of ATP homeostasis in the cardiac muscle, or BIOMD433, a model of MAPK signalling robustness, or BIOMD355, a model of calcium homeostasis, were tested to have homeostatic species.

The results of the ACR test can be interpreted along the same line. Among the 33 models tested, only 3 models passed positively the ACR test. Among these, BIOMD413 verifies the conditions of the Shinar-Feinberg theorem [Shinar and Feinberg, 2010], but the other two models (BIOMD489 and BIOMD738) do not since the deficiency of their reaction network is different from 1. This confirms that these conditions are sufficient, but not necessary. Our new examples could be the starting point of research on more general conditions for ACR.

Finally, two low complexity models, BIOMD614 and BIOMD629, were pin-pointed by our homeostasis tests. We briefly discuss them here because simple models could unravel generic homeostatic mechanism (for details, see Appendix 8). BIOMD614 is a univariate model describing the irreversible reaction kinetics of the conformational transition of a human hormone, where the variable is the fraction of molecules having undergone the transition, which is inevitably equal to one and thus homeostatic at the steady state [Kamihira et al., 2000]. BIOMD629 has 2 reactions and 5 species and the homeostasis found corresponds to buffering, a homeostasis mechanism well known in biochemistry: a buffer is a molecule occurring in much larger amounts than its interactors and whose concentration is practically constant.

For future work, several directions will be investigated. Homeostasis bench-marking was restricted to non-oscillating steady states to avoid a detection of a second steady state inside the oscillatory component, due to equality relaxation used in `IbexOpt`, that could break the homeostasis test. This includes limit cycles and foci. However, oscillations are ubiquitous in biology and it is worth extending our homeostasis definitions to these cases as well.

Other possible improvements concern the performances. As one could see, many models tested cannot be solved within the timeout. If we can explain this with a high dimension of the model, several improvements are possible. First, we know that some convexification contractors using affine arithmetic [Messine and Touhami, 2006; Messine, 2002], or specific to quadratic forms, in particular bi-linear forms [McCormick, 1976] (and the pattern *x*_1_ *x*_2_ is often present in chemical reaction networks) could improve the contraction part of our strategy.

Another promising improvement should be to exploit that these models come from ODEs. Indeed, if a steady state is attractive, it is possible to perform a simulation (with fixed parameters) starting from a random or chosen point of the box that can end near this steady state. Then, a hybrid strategy using this new simulation and the branch and bound strategy should be able to get a feasible point more easily, rendering the comparison for homeostasis more efficient. Moreover this simulation could be used to check if a steady state found is unstable or not. Also, adding as a stop criterion the homeostasis test could be another way to improve it, instead of checking it at a given time.

## 8 Acknowledgements

We thank M.Golubiski and F.Antoneli for presenting us the problem of homeostasy, U.Bhalla and N.Ramakrishnan for sharing their data, C.Lüders for parsing Biomodels, and F.Fages, G. Chabert and B. Neveu for useful discussions. The Ibex computations were performed on the high performance facility MESO@LR of the University of Montpellier. This work is supported by the PRCI ANR/DFG project Symbiont.

## Appendix A1 homeostasis of low complexity models

BIOMD614 is a one species model, with equation:

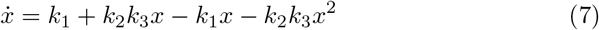

At steady state, this leads to:

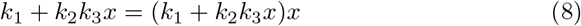

If *k*_1_ ≠ 0, the only solution to (8) is *x* = 1, which is the answer given by IbexHomeo. The model describes the irreversible reaction kinetics of the conformational transition of a human hormone, where *x* is the fraction of molecules having undergone the transition, which is inevitably equal to one at the steady state [Kamihira et al., 2000].

BIOMD629 has 2 reactions and 5 species, provides a 2-homeostasis for kinetics parameters with conserved total amounts fixed, provided by the SBML file. This model does not provide ACR, and the homeostasis found can be explained by the conserved total amounts, that lock species to a small interval. But if we change these total amounts and try again an homeostasis test, it should fail. Indeed this model is given by the equations :

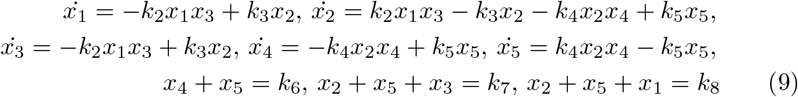

Here *x*_3_ (receptor), and *x*_4_ (coactivator) have been found homeostatic w.r.t. variations of the kinetics parameters. The total amounts are *k*_6_ = 30, 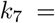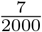, 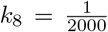. With these values, we get 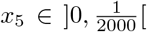, which implies *x*_4_ ∈]29.9995, 30.0005[. In the same way we have 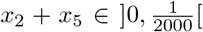, which implies 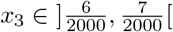. If *k*_6_*, k*_7_*, k*_8_ were closer to each other, there would be no reason for homeostasis. This example corresponds to the homeostasis mechanism known in biochemistry as buffering: a buffer is a molecule in much larger amounts than its interactors and whose concentration is practically constant.

## Appendix A2 Multistationarity statistics

Among the 297 models tested for multistationarity using `IbexSolve`, 63 provide a timeout. For the solved models, 35 do not have steady-state, 153 have a unique steady state, 42 provide multistationarity, 4 have a continuum of steady states. Theses results are given by `IbexSolve`, but some multistationarity models, such as 003, have in reality an oscillatory behavior.

**Table 4.** Time statistics for multistationarity

| # Models | size (min/max/median/average) | median time (s) |
| --- | --- | --- |
| 63 (timeout) | 3/194/23/37.26 | 10800 |
| 234 (solved) | 1/166/8/13.89 | 0.009439 |

Multistationnary models :

- 10 steady states : 703.
- 7 steady states : 435.
- 4 steady states : 228, 249, 294, 517, 518, 663, 709.
- 3 steady states : 003, 008, 026, 027, 029, 069, 116, 166, 204, 233, 257, 296, 447, 519, 573, 625, 630, 687, 707, 708, 714, 729.
- 2 steady states : 079, 100, 156, 230, 315, 545, 552, 553, 688, 713, 716.

## Appendix A3 Homeostasis results tables

**Table 5.**
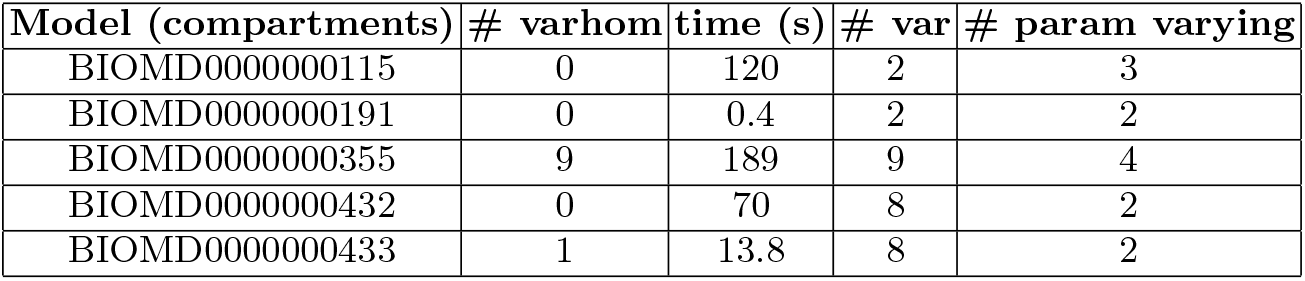
Models with less than 9 species tested for homeostasis w.r.t. the volume of compartments. Minimum and maximum volume of each compartment is set as value given by the SBML file divided by 10 and multiplied by 10, respectively. Kinetic rates and total amount of conservation laws stay fixed. In BIOMD355, all species are 2-homeostatic (and seems independent of the volume). In BIOMD433, the species MK P is 2-homeostatic.

| Model (compartments) | # varhom | time (s) | # var | # param varying |
| --- | --- | --- | --- | --- |
| BIOMD0000000115 | 0 | 120 | 2 | 3 |
| BIOMD0000000191 | 0 | 0.4 | 2 | 2 |
| BIOMD0000000355 | 9 | 189 | 9 | 4 |
| BIOMD0000000432 | 0 | 70 | 8 | 2 |
| BIOMD0000000433 | 1 | 13.8 | 8 | 2 |

**Table 6.** Models with less than 11 species tested for ACR, the second number indicate the number of species that verify 2-homeostasis (ACR included). The minimum and maximum value of each total amount of conservation laws is the value computed from the SBML file, divided by 10 and multiplied by 10, respectively. Kinetic rates and volume compartments stay fixed. FeinbergShinar models serve as tests to confirm that we detect ACR. In BIOMD041, ATP and ATPi are detected to be 2-homeostatic despite the timeout. In BIOMD622, R1B, R1Bubd, and Z are also detected to be 2-homeostatic despite the timeout. In BIOMD413, auxin verify ACR. In BIOMD738, FeDuo, FeRBC, FeSpleen, FeLiver, Hepcidin, FeRest, FeBM are ACR.

| Model (for ACR) | # ACR-# hom | time (s) | # var | # param varying |
| --- | --- | --- | --- | --- |
| BIOMD0000000031 | 0-0 | 3.81 | 3 | 1 |
| BIOMD0000000041 | ?-$\geq 2$ | timeout (2830) | 10 | 3 |
| BIOMD0000000057 | 0-0 | 0.56 | 6 | 1 |
| BIOMD0000000060 | 0-0 | 0.19 | 4 | 1 |
| BIOMD0000000084 | 0-0 | 2.39 | 8 | 4 |
| BIOMD0000000150 | 0-0 | 1.13 | 4 | 2 |
| BIOMD0000000213 |  | timeout (1987) | 6 | 1 |
| BIOMD0000000258 | 0-0 | 3.15 | 3 | 1 |
| BIOMD0000000405 | 0-0 | 0.52 | 5 | 1 |
| BIOMD0000000413 | 1-1 | 6.68 | 5 | 1 |
| BIOMD0000000423 |  | timeout (3071) | 9 | 3 |
| BIOMD0000000432 |  | timeout (2711) | 8 | 3 |
| BIOMD0000000433 |  | timeout (2709) | 8 | 3 |
| BIOMD0000000454 | 0-0 | 0.3 | 3 | 1 |
| BIOMD0000000622 | ?-$\geq 3$ | timeout (3606) | 11 | 1 |
| BIOMD0000000629 | 0-0 | 4.32 | 5 | 3 |
| BIOMD0000000646 | 0-0 | 801 | 11 | 1 |
| BIOMD0000000647 | 0-0 | 306 | 11 | 5 |
| BIOMD0000000738 | 7-7 | 557 | 11 | 1 |
| FeinbergShinar1 | 1-1 | 5.31 | 7 | 2 |
| FeinbergShinar2 | 1-1 | 3.08 | 8 | 2 |
| FeinbergShinar3 | 1-1 | 15.23 | 5 | 2 |

**Table 7.** Models with less than 9 species tested for homeostasis w.r.t. kinetics rates. The minimum and maximum value of each kinetic rate is given by the SBML file divided by 100 and multiplied by 100, respectively. In BIOMD614, the unique species is 2-homeostatic (moreover independent). In BIOMD629, receptor and coactivator are 2-homeostatic.

| Models (kinetics) | # varhom | time (s) | # var | # param varying |
| --- | --- | --- | --- | --- |
| BIOMD0000000023 | 0 | 301 | 5 | 60 |
| BIOMD0000000031 | 0 | 93.8 | 3 | 12 |
| BIOMD0000000057 | 0 | 284 | 6 | 25 |
| BIOMD0000000060 | 0 | 122 | 4 | 10 |
| BIOMD0000000065 |  | timeout (2885) | 8 | 23 |
| BIOMD0000000067 |  | timeout (2523) | 7 | 20 |
| BIOMD0000000076 | 0 | 60 | 1 | 20 |
| BIOMD0000000084 | 0 | 22 | 8 | 17 |
| BIOMD0000000115 |  | crashed | 2 | 7 |
| BIOMD0000000150 | 0 | 243 | 4 | 6 |
| BIOMD0000000159 | 0 | 91 | 3 | 6 |
| BIOMD0000000191 | 0 | 89 | 2 | 19 |
| BIOMD0000000203 | 0 | 876 | 5 | 34 |
| BIOMD0000000213 |  | timeout (2173) | 6 | 46 |
| BIOMD0000000219 | 0 | 540 | 9 | 74 |
| BIOMD0000000221 |  | timeout (2902) | 8 | 53 |
| BIOMD0000000222 |  | timeout (2909) | 8 | 53 |
| BIOMD0000000228 | 0 | 1147 | 9 | 40 |
| BIOMD0000000240 |  | timeout (2163) | 6 | 20 |
| BIOMD0000000249 |  | timeout (2883) | 8 | 7 |
| BIOMD0000000258 | 0 | 132 | 3 | 9 |
| BIOMD0000000284 | 0 | 31 | 6 | 3 |
| BIOMD0000000325 | 0 | 300 | 5 | 16 |
| BIOMD0000000355 |  | timeout (2287) | 9 | 22 |
| BIOMD0000000405 | 0 | 801 | 5 | 5 |
| BIOMD0000000413 | 0 | 481 | 5 | 12 |
| BIOMD0000000414 | 0 | 3.7 | 1 | 4 |
| BIOMD0000000417 |  | timeout (360) | 1 | 12 |
| BIOMD0000000423 |  | timeout (3132) | 9 | 16 |
| BIOMD0000000425 | 0 | 0.4 | 1 | 6 |
| BIOMD0000000432 | $\leq 1$ | timeout (1031) | 8 | 26 |
| BIOMD0000000433 | $\leq 3$ | timeout (1671) | 8 | 26 |
| BIOMD0000000454 |  | timeout (1022) | 3 | 11 |
| BIOMD0000000456 |  | timeout (1375) | 4 | 16 |
| BIOMD0000000458 | 0 | 93 | 2 | 15 |
| BIOMD0000000459 | 0 | 63 | 3 | 8 |
| BIOMD0000000460 | 0 | 122 | 3 | 8 |
| BIOMD0000000495 | 0 | 365 | 9 | 36 |
| BIOMD0000000519 |  | timeout (1050) | 3 | 8 |
| BIOMD0000000530 | 0 | 1923 | 7 | 17 |
| BIOMD0000000590 | 0 | 540 | 9 | 30 |
| BIOMD0000000614 | 1 | 17 | 1 | 3 |
| BIOMD0000000615 | 0 | 160 | 4 | 12 |
| BIOMD0000000626 |  | timeout (632) | 6 | 20 |
| BIOMD0000000629 | 2 | 1 | 5 | 4 |
| BIOMD0000000708 | 0 | 1770 | 5 | 13 |
| BIOMD0000000728 | 0 | 1 | 2 | 4 |

**Table 8.** Models with more than 10 species tested for homeostasis w.r.t. the volume of compartments. Minimum and maximum volume of each compartment is set as value given by the SBML file divided by 10 and multiplied by 10, respectively. Kinetic rates and total amount of conservation laws stay fixed. In BIOMD738, we know that at least Hepcidin is 2-homeostatic (and independent) despite the timeout.

| Model (compartments) | # varhom | time (s) | # var | # param varying |
| --- | --- | --- | --- | --- |
| BIOMD0000000041 |  | timeout (3759) | 10 | 2 |
| BIOMD0000000093 |  | timeout (12901) | 34 | 2 |
| BIOMD0000000123 |  | timeout (5681) | 14 | 2 |
| BIOMD0000000192 |  | timeout (3924) | 13 | 2 |
| BIOMD0000000482 |  | timeout (8308) | 23 | 3 |
| BIOMD0000000491 |  | timeout (20962) | 57 | 3 |
| BIOMD0000000492 |  | timeout (19066) | 52 | 3 |
| BIOMD0000000581 |  | timeout (5359) | 27 | 2 |
| BIOMD0000000738 | $\geq 1$ | timeout (3676) | 11 | 7 |

**Table 9.** Models with more than 12 species tested for ACR, the second number indicate the number of species that verify 2-homeostasis (ACR included). The minimum and maximum value of each total amount of conservation laws is the value computed from the SBML file, divided by 10 and multiplied by 10, respectively. Kinetic rates and volume compartments stay fixed. In BIOMD489, LPS:LBP:CD14:TLR4:TIRAP:MyD88:IRAK4, IkBb mRNA, IkBe mRNA, LPS:LBP:CD14:TLR4:RIP1:TRAM:TRIF:TBK/IKKe are detected ACR despite the timeout.

| Model (for ACR) | # ACR-# hom | time (s) | # var | # param varying |
| --- | --- | --- | --- | --- |
| BIOMD0000000009 |  | timeout (7988) | 22 | 7 |
| BIOMD0000000011 |  | timeout (7804) | 22 | 7 |
| BIOMD0000000030 |  | timeout (6318) | 18 | 3 |
| BIOMD0000000038 |  | timeout (4400) | 13 | 4 |
| BIOMD0000000048 |  | timeout (8380) | 23 | 6 |
| BIOMD0000000093 |  | timeout (12115) | 34 | 6 |
| BIOMD0000000123 | 0-0 | 4810 | 14 | 3 |
| BIOMD0000000192 | 0-0 | 1316 | 13 | 3 |
| BIOMD0000000270 | 0-0 | 10887 | 32 | 9 |
| BIOMD0000000431 |  | timeout (9808) | 27 | 6 |
| BIOMD0000000489 | $\geq 4 - \geq 4$ | timeout (11104) | 52 | 4 |
| BIOMD0000000491 |  | timeout (19323) | 57 | 1 |
| BIOMD0000000492 |  | timeout (17524) | 52 | 1 |
| BIOMD0000000581 |  | timeout (8485) | 27 | 10 |

**Table 10.** Models with more than 10 species tested for homeostasis w.r.t. kinetics rates. The minimum and maximum value of each kinetic rate is given by the SBML file divided by 100 and multiplied by 100, respectively. In BIOMD048, EGF is detected 2-homeostatic despite the timeout. In BIOMD093, SHP2 is detected homeostatic despite the crash.

| Models (kinetics) | # varhom | time (s) | # var | # param varying |
| --- | --- | --- | --- | --- |
| BIOMD0000000009 |  | timeout (8239) | 22 | 32 |
| BIOMD0000000011 |  | timeout (8009) | 22 | 30 |
| BIOMD0000000028 |  | timeout (5810) | 16 | 27 |
| BIOMD0000000030 |  | timeout (6571) | 18 | 32 |
| BIOMD0000000038 |  | timeout (4722) | 13 | 24 |
| BIOMD0000000041 |  | timeout (3952) | 10 | 25 |
| BIOMD0000000048 | $\geq 1$ | timeout (9551) | 23 | 50 |
| BIOMD0000000093 | $\geq 1$ | crashed | 34 | 73 |
| BIOMD0000000123 |  | timeout (5654) | 14 | 22 |
| BIOMD0000000192 |  | timeout (3923) | 13 | 18 |
| BIOMD0000000218 | 0 | 541 | 12 | 70 |
| BIOMD0000000270 |  | timeout (12271) | 32 | 30 |
| BIOMD0000000294 |  | timeout (3608) | 10 | 10 |
| BIOMD0000000388 |  | timeout (1801) | 11 | 19 |
| BIOMD0000000431 |  | timeout (10028) | 27 | 44 |
| BIOMD0000000482 |  | timeout (8452) | 23 | 56 |
| BIOMD0000000489 |  | timeout (15513) | 52 | 105 |
| BIOMD0000000491 |  | timeout | 57 | 172 |
| BIOMD0000000492 |  | timeout | 52 | 176 |
| BIOMD0000000581 |  | timeout (9131) | 27 | 35 |
| BIOMD0000000622 |  | timeout (3618) | 11 | 25 |
| BIOMD0000000646 |  | timeout (3976) | 11 | 33 |
| BIOMD0000000647 |  | timeout (1173) | 11 | 11 |
| BIOMD0000000707 |  | timeout (1746) | 5 | 10 |
| BIOMD0000000738 |  | timeout (3986) | 11 | 33 |

